## supplemental figures for "Control of meiotic entry by dual inhibition of a key mitotic transcription factor"

Figure 1 Supplement

### Su et al_Figure S1

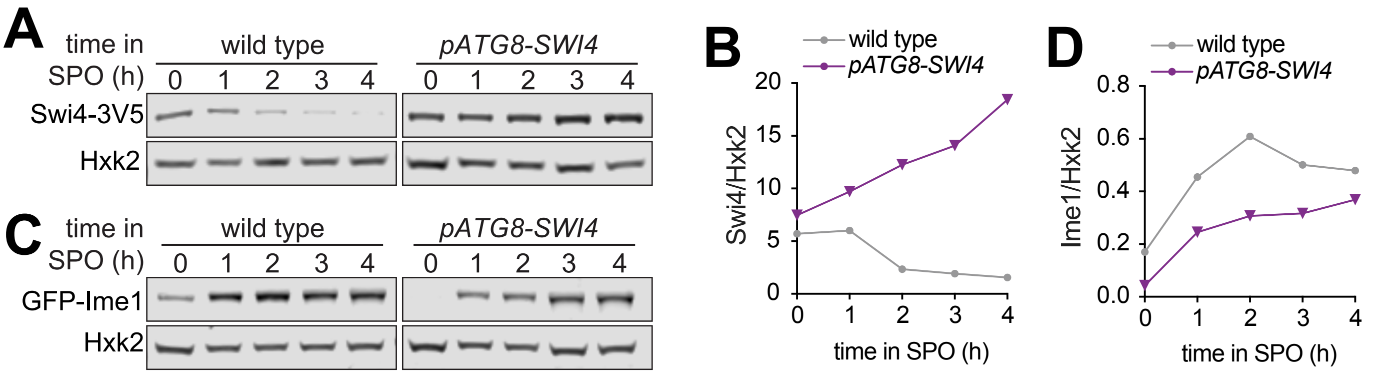

Figure 2 Supplement

### Su et al_Figure S2

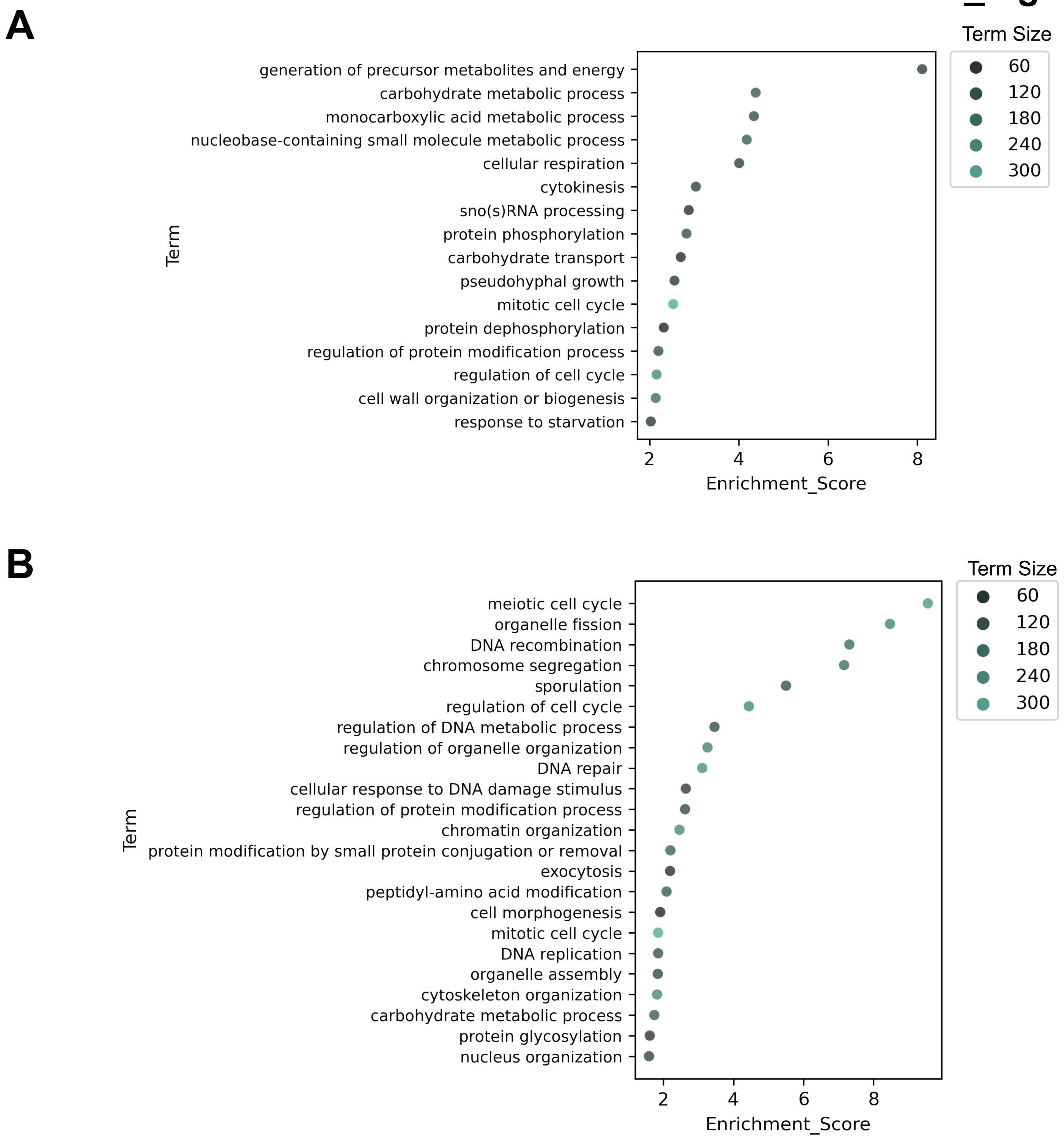

### Su et al_Figure S3

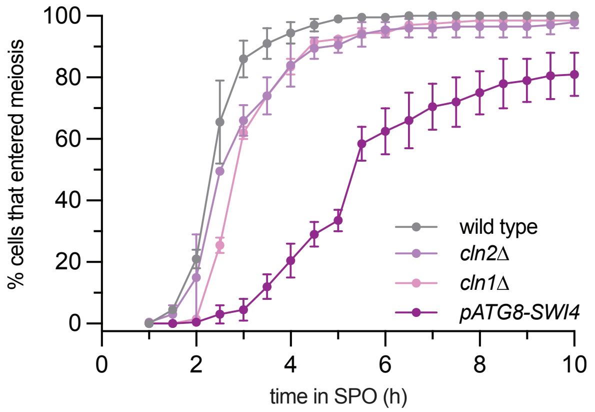

Figure 4 Supplement

**Su et al_Figure S4**

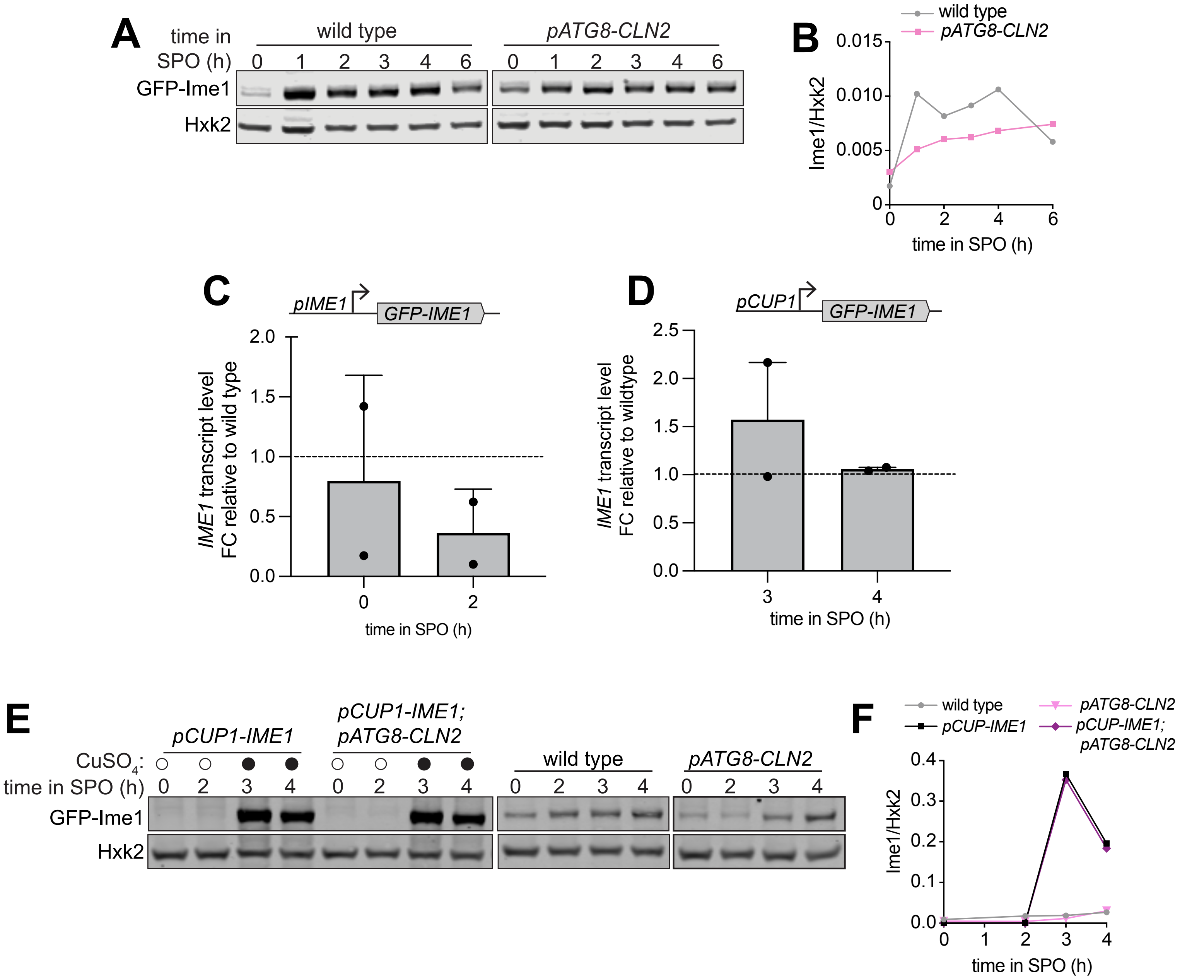

### Su et al_Figure S5

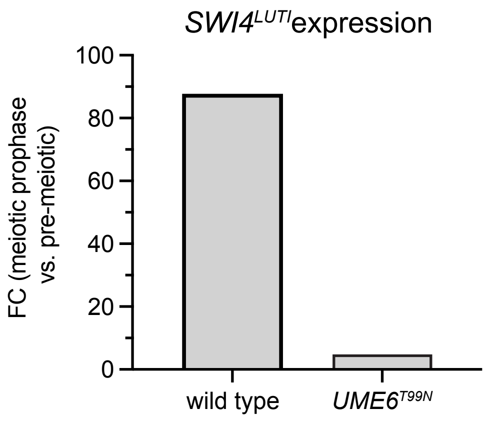

Figure 6 Supplement

### Su et al_Figure S6

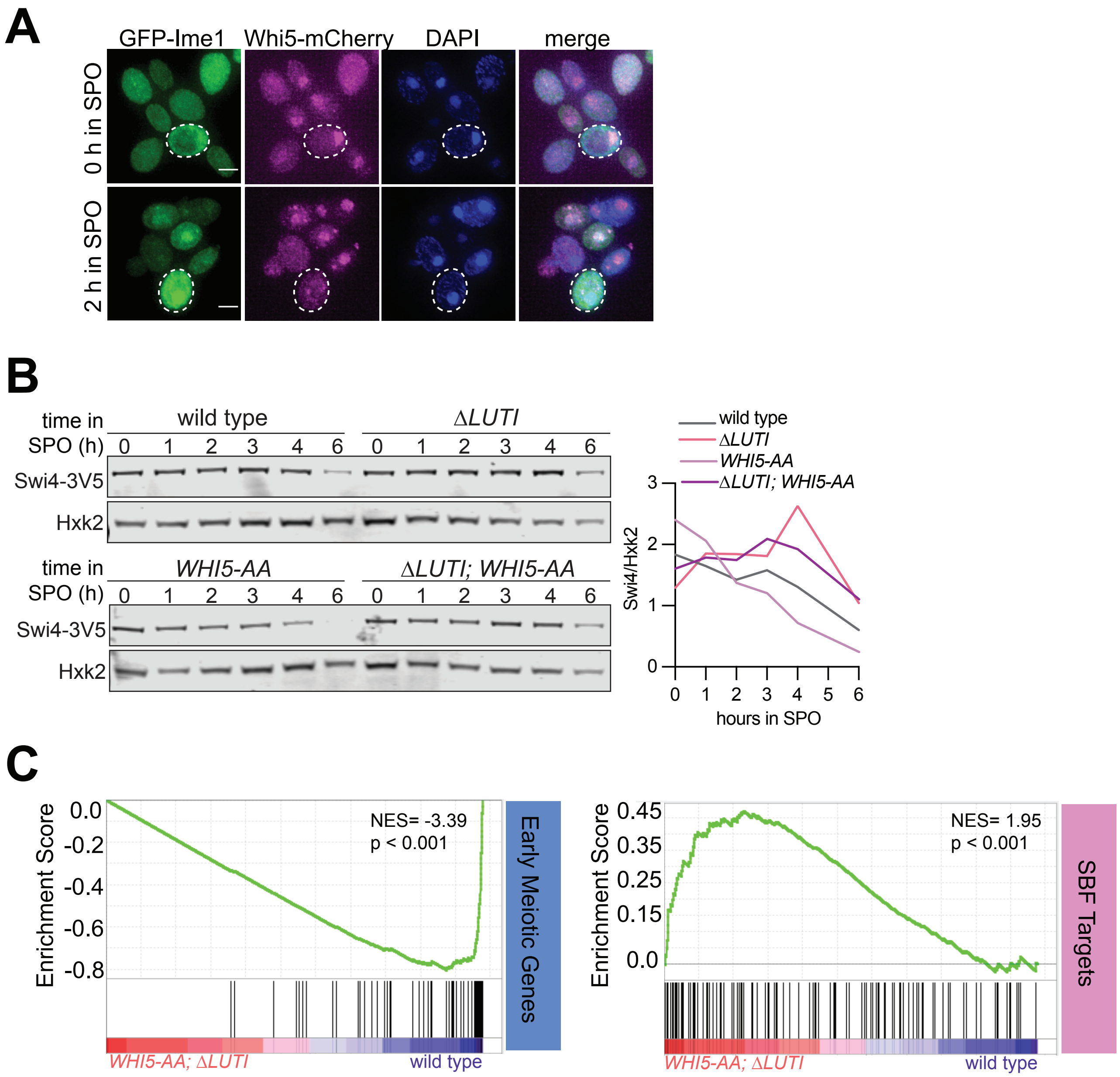
